# USP30 inhibition induces mitophagy and reduces oxidative stress in parkin-deficient and CCCP-stressed human iPSC-derived neurons

**DOI:** 10.1101/2022.06.29.498100

**Authors:** Justyna Okarmus, Jette Bach Agergaard, Tina C. Stummann, Henriette Haukedal, Malene Ambjørn, Kristine K. Freude, Karina Fog, Morten Meyer

**Affiliations:** Department of Neurobiology Research, Institute of Molecular Medicine, University of Southern Denmark, J.B. Winsløws Vej 21, 5000 Odense C, Denmark; Neuroscience, H. Lundbeck A/S, Ottiliavej 9, 2500 Valby, Denmark; Department of Veterinary and Animal Sciences, Faculty of Health and Medical Sciences, University of Copenhagen, Grønnegaardsvej 7, 1870 Frederiksberg C, Denmark; Department of Neurology, Odense University Hospital, J.B. Winsløws Vej 4, 5000 Odense C, Denmark; BRIDGE - Brain Research Inter-Disciplinary Guided Excellence, Department of Clinical Research, University of Southern Denmark, J.B. Winsløws Vej 19, 5000 Odense C, Denmark

**Author notes:** Corresponding author: Morten Meyer, Ph.D. Department of Neurobiology Research Institute of Molecular Medicine University of Southern Denmark J.B. Winsløws Vej 21, st. DK-5000 Odense C, Denmark.

**Keywords:** parkin, mitochondria, dopamine, ROS production, induced pluripotent stem cells, Parkinson’s disease

## Abstract

Ubiquitination of mitochondrial proteins plays an important role in the cellular regulation of mitophagy. The E3 ubiquitin ligase parkin (encoded by *PARK2*) and the ubiquitin-specific protease 30 (USP30) have both been reported to regulate ubiquitination of outer mitochondrial proteins and thereby mitophagy. Loss of E3 ligase activity is thought to be pathogenic in both sporadic and inherited Parkinson’s disease (PD), with loss-of-function mutations in *PARK2* being the most frequent cause of autosomal recessive PD.

The aim of the present study was to evaluate whether mitophagy induced by USP30 inhibition provides a functional rescue in isogenic human induced pluripotent stem cell-derived dopaminergic neurons with and without *PARK2* knockout (KO).

Our data show that healthy neurons responded to CCCP-induced mitochondrial damage by clearing the impaired mitochondria and that this process was accelerated by USP30 inhibition. Parkin-deficient neurons showed an impaired mitophagic response to CCCP challenge, although mitochondrial ubiquitination was enhanced. USP30 inhibition promoted mitophagy in *PARK2* KO neurons, independently of whether left in basal conditions or treated with CCCP. In *PARK2* KO, as in control neurons, USP30 inhibition balanced oxidative stress levels by reducing excessive production of reactive oxygen species. Interestingly, non-dopaminergic neurons, were the main driver of the beneficial effects of USP30 inhibition.

Our findings demonstrate that USP30 inhibition is a promising approach to boost mitophagy and improve cellular health, also in parkin-deficient cells, and support the potential relevance of USP30 inhibitors as a novel therapeutic approach in diseases with a need to combat neuronal stress mediated by impaired mitochondria.

## Introduction

Parkinson’s disease (PD) is a neurodegenerative movement disorder characterized by the progressive loss of mainly midbrain dopaminergic neurons [1]. Despite years of research, there is no cure for PD. Current treatments are symptomatic only and based on dopaminergic drugs that are associated with dyskinesia and reduced effect as the disease progresses [2–4]. Development of alternative PD treatment strategies is therefore of high interest.

The most frequent cause of autosomal recessive PD is loss-of-function mutations in the *PARK2* gene, which encodes for the protein parkin [5]. Parkin inactivation has also been suggested to be implicated in sporadic PD [6, 7]. Investigations of cellular effects of mutations in the *PARK2* gene may therefore reveal common disease mechanisms important for unraveling the pathogenesis of both familial and sporadic PD and thereby guide identification of novel therapeutic strategies.

Parkin is a key player in regulation of mitophagy [8]. Loss of the mitochondrial membrane electrochemical potential leads to dysfunctional oxidative phosphorylation. The membrane depolarization is followed by stabilization of phosphatase and tensin homolog induced kinase 1 (PINK1) at the mitochondria membrane, where it phosphorylates and recruits parkin to the mitochondria [9]. Parkin recruitment leads to increased ubiquitination of outer mitochondrial membrane proteins, which is further stabilized by PINK1 phosphorylation of ubiquitin at serine 65 (pS65-Ub). High total ubiquitination load is the signal for engulfment of the mitochondria by the autophagosome [10, 11]. Mitophagy protects against the accumulation of damaged mitochondria and toxic reactive oxygen species (ROS) [12] and is therefore expected to be neuroprotective [13].

Ubiquitination and deubiquitination are enzymatically mediated processes by which ubiquitin is covalently bound or cleaved from proteins by E3 ubiquitin ligases or deubiquitinating enzymes, respectively [14, 15]. USP30, a deubiquitinase, has been reported to counteract parkin-dependent mitophagy by deubiquitinating outer mitochondrial membrane proteins, in particular TOM20 [13, 16]. Knockdown (KD) of USP30 in parkin-overexpressing cells promotes the clearance of mitochondria in response to mitochondrial depolarizing agents [17]. This suggests that USP30 inhibition may be a therapeutic approach to facilitate mitophagy of dysfunctional mitochondria and rescue neurons.

USP30 KD or inhibition can induce mitophagy in cultured cells, including murine primary neurons [13,18,19]. USP30 inhibition was recently shown to increase cardiac mitophagy *in vivo* [19]. Less is known about the relevance of USP30 in controlling mitophagy induction in the human brain, and specifically in brains of *PARK2* mutation carriers. Induced pluripotent stem cells (iPSCs) allow for differentiation into human midbrain dopaminergic neurons [20, 21]. In this study, we took advantage of the iPSC technology to generate isogenic human dopaminergic neurons with and without *PARK2* knockout (KO). This enabled us to study the effect of *PARK2* KO on induction of mitophagy upon chemical mitochondrial damage and the potential of USP30 inhibition to boost mitophagy and decrease oxidative stress in neurons with and without functional parkin protein.

## Results

### Isogenic *PARK2* KO and control cultures contain similar cell populations

To study the disease mechanisms underlining *PARK2*-mediated PD, we compared healthy human iPSCs and isogenic iPSCs where the PARK2 gene was deleted using zinc finger nuclease gene editing [22]. Detailed information on this cell line can be found in our recent studies [23–26]. Isogenic *PARK2* KO and control iPSC-derived neural stem cells (NSCs) were differentiated into neuronal cultures (*Fig. 1*) containing a significant proportion of midbrain dopaminergic neurons as previously demonstrated [23, 25]. The neurons had defined long neurites and expressed beta-TubulinIII (a marker of newly formed neurons), MAP2 (a marker of mature neurons), and synaptophysin (a marker of presynaptic regions). As expected, many of the neurons (approximately 35%) were also positive for tyrosine hydroxylase (TH), indicating dopaminergic identity, and FOXA2, indicating a midbrain phenotype. Moreover, the cultures contained a population of GABA+ neurons and GFAP+ astrocytes. The differentiation of the isogenic cell lines resulted in neuronal cultures with similar morphologies and distribution of cellular subtypes.

**Figure 1:**
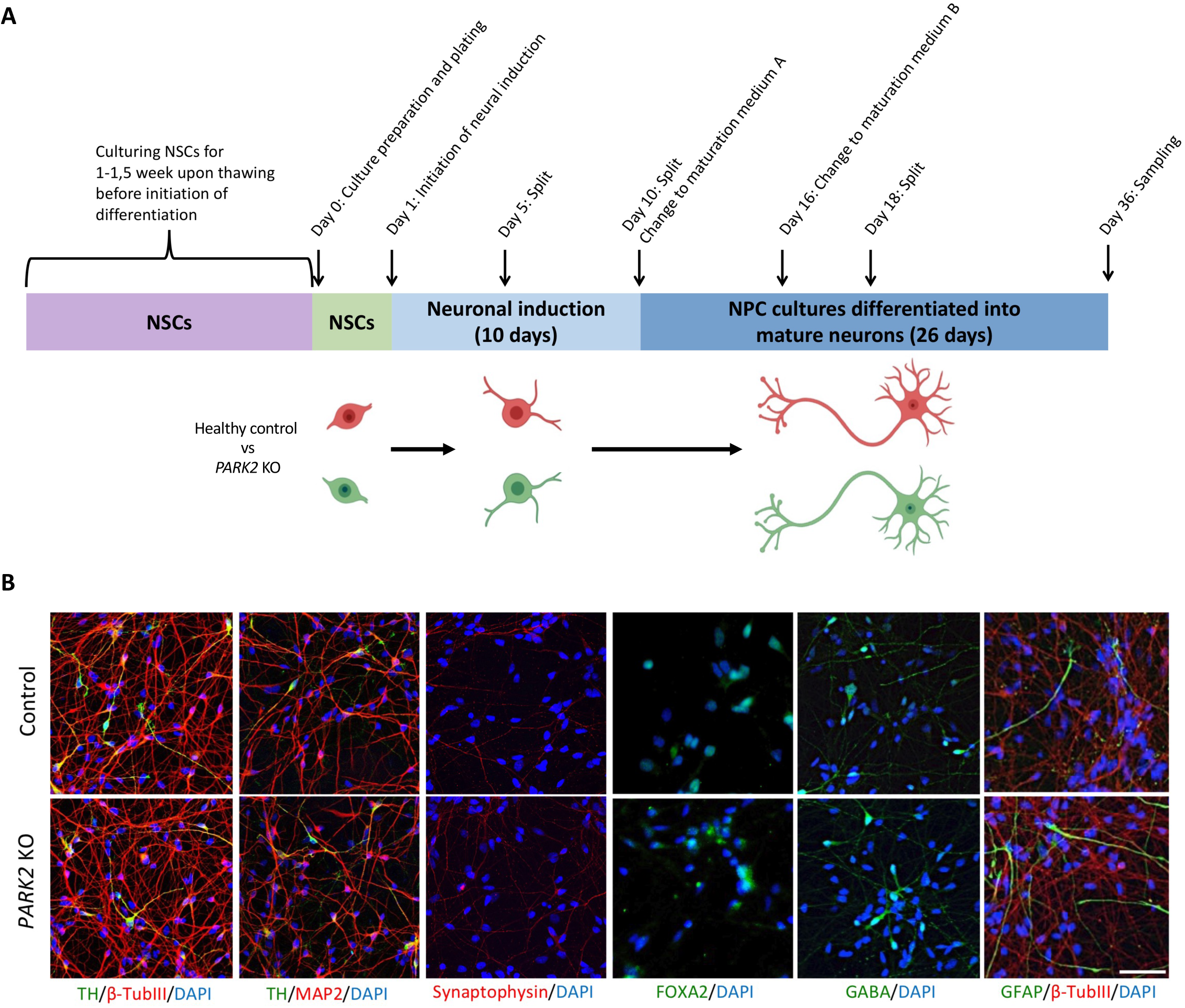
Derivation and characterization of healthy and *PARK2* KO neurons. **A)** Graphical overview of the differentiation of isogenic iPSC-derived neural stem cells (NSCs) into fully committed mature neurons. **B)** Immunofluorescence staining of control and *PARK2* KO NSCs differentiated into mature cells showed a large proportion of beta-TubulinIII+ newly formed neurons (red), MAP2+ mature neurons (red), including a significant population of tyrosine hydroxylase (TH)+ dopaminergic neurons (green). The cells also expressed synaptophysin (red, a marker of presynaptic regions) and FOXA2 (green, a marker of the midbrain phenotype). A small population of GABA+ neurons and GFAP+ astrocytes was also present in the cultures. Cell nuclei were stained with DAPI (blue). Scale bar = 50 μm.

### Parkin deficiency results in impaired mitophagy

We have previously demonstrated increased mitochondrial area and accumulation of mitochondria with abnormal morphology in *PARK2* KO neurons using transmission electron microscopy (TEM) [26]. The current study confirmed the increased mitochondrial area (*Figs. 2 AB*) in *PARK2* KO neurons compared to that of control neurons at basal growth conditions (no carbonyl cyanide m-chlorophenylhydrazone (CCCP)-mediated mitochondrial stress) using image quantification of immunofluorescent staining for the mitochondrial markers TOM20 and HSP60. Supporting this, Western blotting for TOM20 and HSP60, and ultrastructural analysis (no CCCP, *Figs. 2 C-F*) showed a non-significant tendency for increased mitochondrial area in *PARK2* KO neurons. The impaired mitophagy in *PARK2* KO neurons was correlated with elevated ROS levels (no CCCP, *Fig. 2G*).

**Figure 2:**
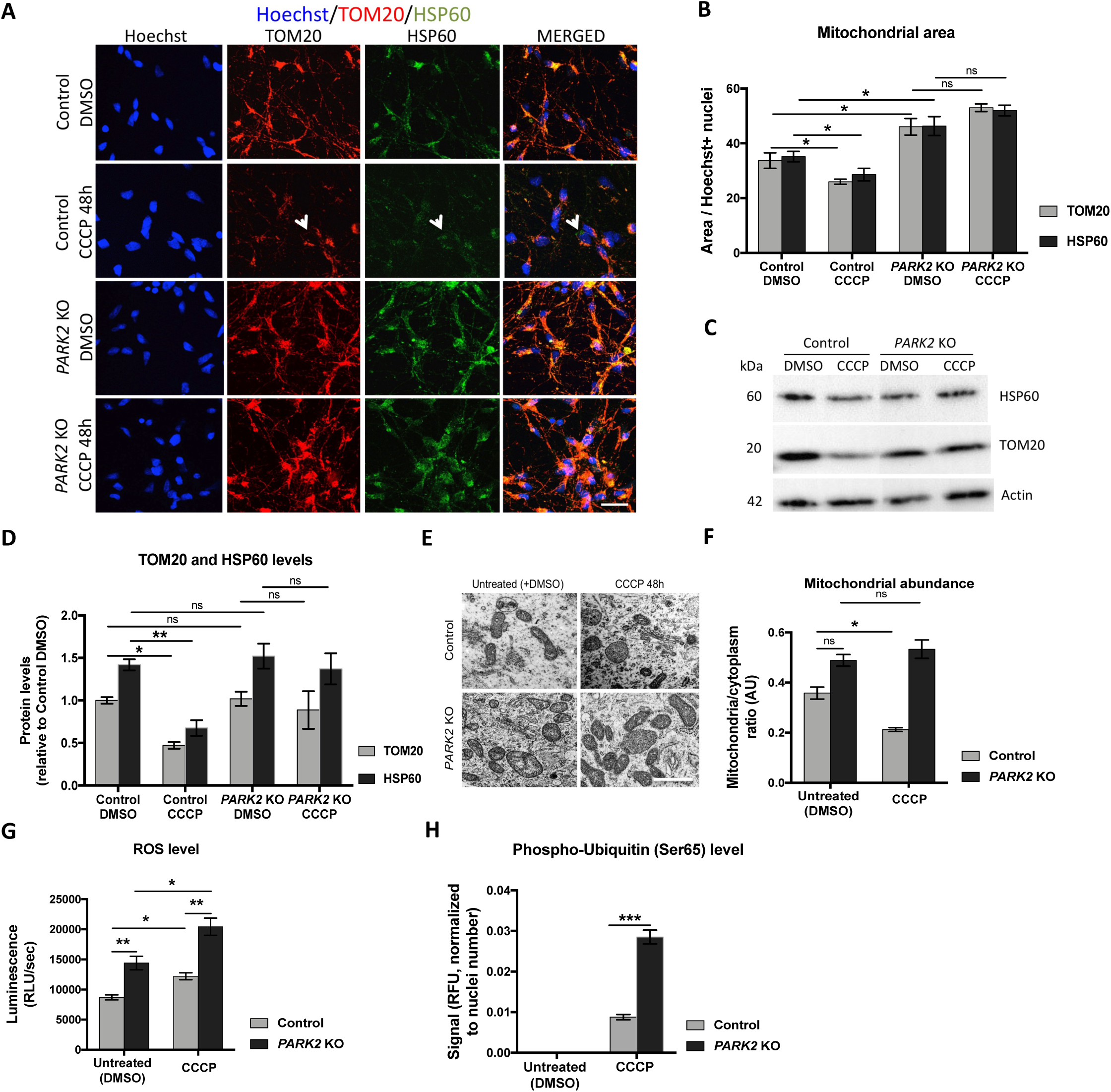
Impaired mitophagy in *PARK2* KO neurons. **A)** Investigation of mitophagy in *PARK2* KO and control neurons after CCCP treatment (10 µM, 48 h) by immunofluorescence staining for the mitochondrial markers TOM20 (red) and HSP60 (green); Hoechst (blue) indicates nuclei. Scale bar: 20 µm. **B)** Quantification of TOM20 and HSP60 immunoreactivity. **C)** Western blotting analysis for TOM20 and HSP60. **D)** Quantification of Western blots. Expression levels were normalized to α-actin and are shown relative to control/DMSO neurons (untreated). **E)** Representative transmission electron microscopy (TEM) micrographs of treated (10 µM CCCP, 48h) and untreated neurons. Scale bar: 500 nm. **F)** Quantification of mitochondria on TEM images in E). **G)** Reactive oxygen species (ROS) levels in control and *PARK2* KO neurons after exposure (10 µM CCCP, 48h). **H)** pS65-Ub was elevated in *PARK2* KO and control neurons after CCCP treatment. pS65-Ub levels were below detection without CCCP. Data presented as mean±SEM, n=3-9 technical replicates, data from 2 (F, H) or 3 (B, D, G) independent differentiations. Significant differences are indicated by *p < 0.05, **p < 0.01, ***p < 0.001, ns: not significant, two-way ANOVA followed by Tukey’s post hoc for multiple comparisons.

To analyze whether the observed mitochondrial accumulation was associated with impaired mitophagy, we monitored the ability of the neurons to clear impaired mitochondria in response to CCCP [27]. To optimize the mitophagy response window, neurons were treated with 10 µM CCCP in a time-dependent manner (24, 48, 72, and 96 hrs) followed by quantification of nuclei and the area covered by the mitochondrial marker TOM20 (*Suppl. Figs. S1A-C*). TOM20 area was unaffected by CCCP treatment in *PARK2* KO neurons. In control neurons, the 48h time point gave the largest reduction in mitochondrial area without affecting cell survival and was used in subsequent experiments.

Studies indicate that the loss of TOM20 staining can reflect proteolysis at the outer mitochondrial membrane by the ubiquitin-proteasome system (UPS) rather than *de facto* degradation of the mitochondria via mitophagy [28, 29]. Therefore, we quantified the area covered by both TOM20 and the mitochondrial matrix protein HSP60 in response to CCCP treatment (*Figs. 2AB*) and showed that both markers decline in response to CCCP treatment in control neurons. Noticeably, the remaining mitochondrial population was positive for both TOM20 and HSP60 and, although we could observe discrete patches negative for TOM20, the TOM20 and HSP60 areas were not significantly different *(Figs. 2AB)*. Western blotting followed by densitometric analysis confirmed the reduction in TOM20 and HSP60 proteins (*Figs. 2CD*), and TEM revealed fewer mitochondria (*Figs. 2EF*) in control neurons in response to CCCP, confirming clearance of CCCP-damaged mitochondria. Mitochondrial load was not decreased by CCCP treatment in *PARK2* KO neurons as indicated by TOM20 and HSP60 immunostaining (*Figs. 2AB*), Western blotting (*Figs. 2CD*), and TEM (*Figs. 2EF*). This implies that parkin deficiency is associated with impaired mitophagy. ROS data showed CCCP-induced mitochondrial stress in both lines, confirming the induction of mitochondrial stress at the 48h time point (*Fig. 2G*).

Following mitochondrial depolarization, PINK1 becomes stabilized on the outer mitochondrial membrane leading to phosphorylation of ubiquitin at serine 65 (pS65-Ub) [30]. Untreated *PARK2* KO and control neurons had undetectable levels of pS65-Ub, which were strongly amplified in both cell lines after CCCP treatment (*Fig. 2H*). The pS65-Ub levels were significantly higher in CCCP-treated parkin-deficient neurons compared to healthy controls (*Fig. 2H*), potentially caused by accumulation of damaged mitochondria in *PARK2* KO neurons.

Overall, our data show that control neurons respond to mitochondrial damage with mitophagy whereas *PARK2* KO neurons are defective in clearing impaired mitochondria.

### USP30 inhibition facilitates mitophagy in control and *PARK2* KO neurons after mitochondrial damage with CCCP and induces basal mitophagy in *PARK2* KO neurons

USP30 counteracts parkin-mediated ubiquitination of mitochondrial outer membrane proteins, thereby inhibiting mitophagy [13, 31] (*Suppl. Fig. S2A*). Therefore, we investigated whether USP30 inhibition could accelerate mitophagy in control and *PARK2* KO neurons after mitochondrial damage with CCCP. Western blotting confirmed that our isogenic cell lines expressed comparable levels of USP30 protein (*Suppl. Fig. S2B*). Release of lactate dehydrogenase (LDH) and counts of cell nuclei (DAPI+ nuclei) identified 3 µM of the USP30 inhibitors USP30i-37 and USP30i-3 in a dose-response experiment to be non-toxic to the neurons, also in the presence of CCCP (*Suppl. Figs. S3, S4*). Moreover, 3 µM of the USP30 inhibitors did not affect the neuronal content in the cultures i.e., no changes were observed in the numbers of MAP2+ and TH+ neurons (*Suppl. Fig. S4)*.

To assess the potential of the compounds to promote mitophagy, neurons were treated with USP30 inhibitors (0-6 µM) for 4 h prior to CCCP exposure (10 µM), and mitophagy was evaluated by double immunofluorescence staining for TOM20 and HSP60 (*Figs. 3A, 4A*). In line with data presented in *Fig. 2B*, CCCP significantly decreased TOM20 and HSP60 areas in healthy control neurons (*Figs. 3A-C*) but did not reduce TOM20 and HSP60 areas in *PARK2* KO neurons (*Figs. 4A-C*).

**Figure 3:**
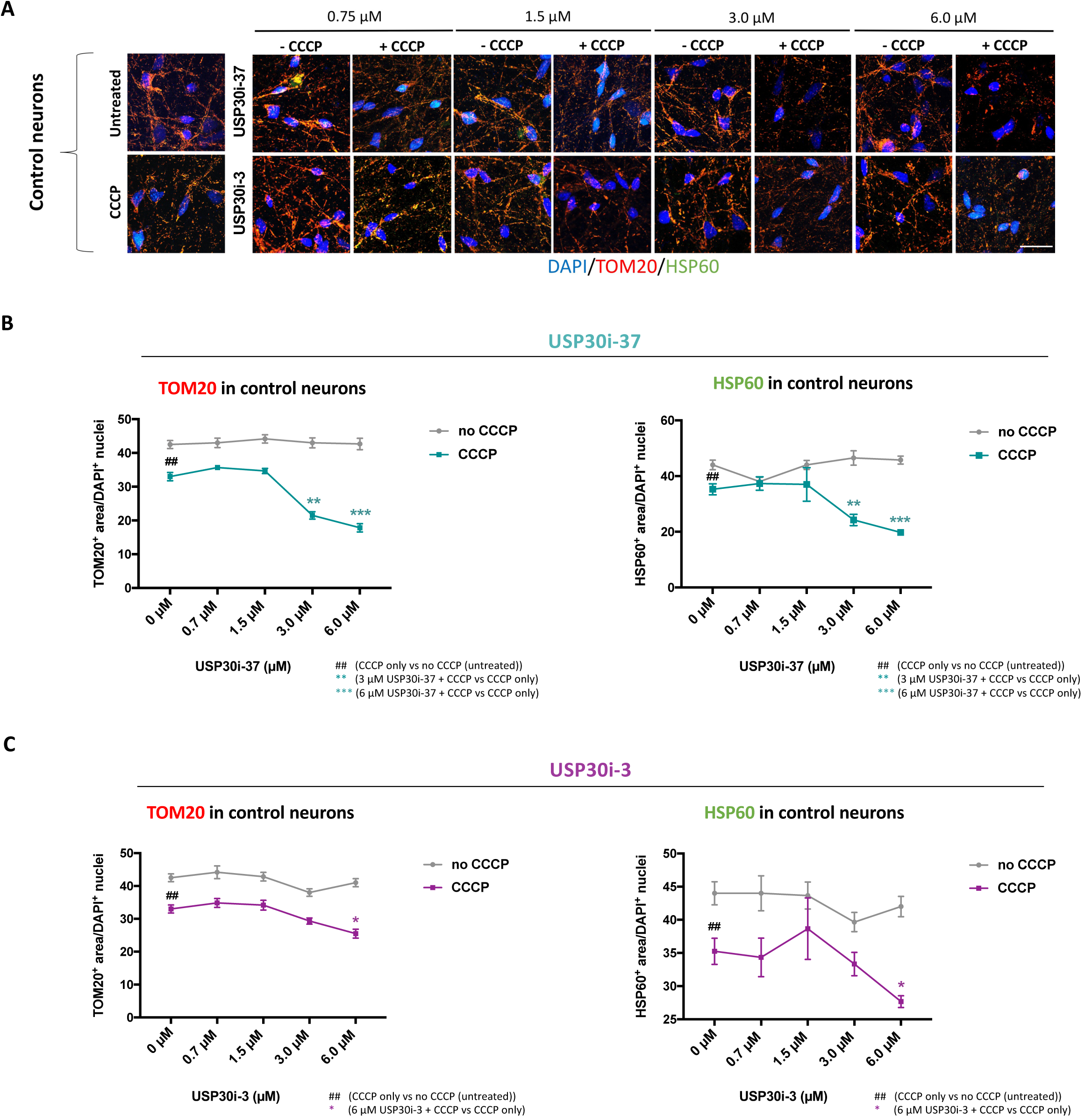
USP30 inhibition enhances CCCP-induced mitophagy but not basal mitophagy in control neurons. **A)** Immunofluorescence staining for the mitochondrial markers TOM20 (red) and HSP60 (green) after treatment with USP30 inhibitors (0.75 µM, 1.5 µM, 3 µM, 6 µM) added 4 h prior to CCCP (10 µM, 48 h). DAPI (blue) indicates nuclei. Scale bar: 50 µm. Quantification of TOM20 and HSP60 immunoreactivity upon treatment with **B)** USP30i-37 and **C)** USP30i-3 alone and in combination with CCCP in control neurons. Data presented as mean±SEM, n=9 technical replicates, data from 3 independent differentiations. Significant differences are indicated by *p < 0.05, **, ^##^p < 0.01, ***p < 0.001, one-way ANOVA followed by Dunnett’s post hoc for multiple comparisons.

**Figure 4:**
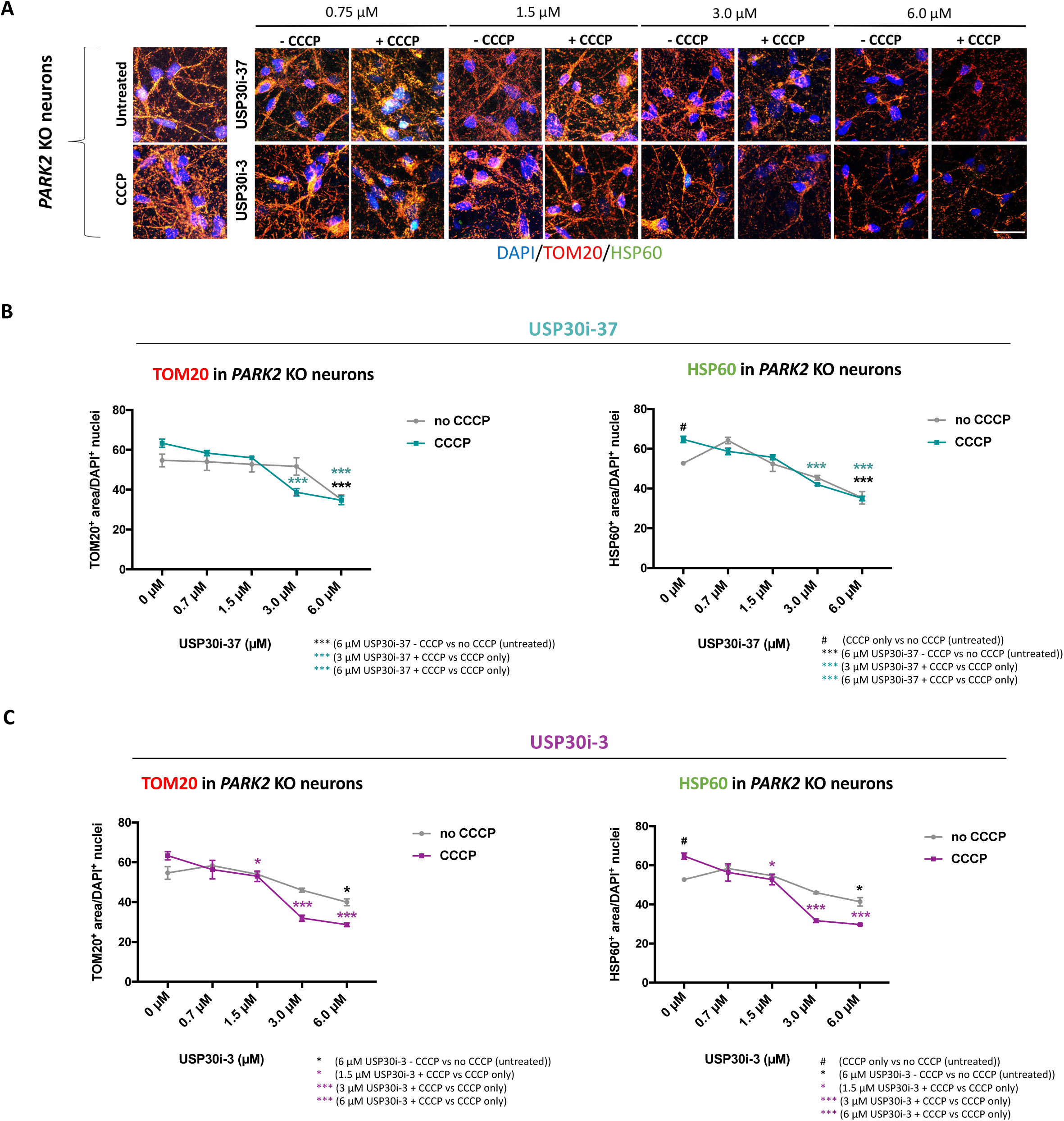
USP30 inhibition induces basal mitophagy and mitophagy of CCCP-damaged mitochondria in *PARK2* KO neurons. **A)** Immunofluorescence staining for the mitochondrial markers TOM20 (red) and HSP60 (green) after treatment with USP30 inhibitors (0.75 µM, 1.5 µM, 3 µM, 6 µM) added 4 h prior to CCCP (10 µM, 48 h). DAPI (blue) indicates nuclei. Scale bar: 50 µm. Quantification of TOM20 and HSP60 immunoreactivity upon treatment with **B)** USP30i-37 and **C)** USP30i-3 alone and in combination with CCCP in *PARK2* KO neurons. Data presented as mean±SEM, n=9 technical replicates, data from 3 independent differentiations. Significant differences are indicated by *, ^#^p < 0.05, ***p < 0.001, one-way ANOVA followed by Dunnett’s post hoc for multiple comparisons.

A significant potentiation of the CCCP-induced decrease in TOM20 and HSP60 areas in control neurons was observed after treatment with 3 µM and 6 µM of USP30i-37 (*Fig. 3B*) and 6 µM of USP30i-3 (*Fig. 3C*), in line with USP30 inhibition boosting mitophagy. Importantly, 3 and 6 µM of both inhibitors boosted the CCCP-induced TOM20 and HSP60 area reduction in the *PARK2* KO neurons as well (*Figs. 4A-C*). This shows that USP30 inhibition can accelerate mitophagy in *PARK2* KO neurons even if mitochondrial damage by CCCP did not do so. In *PARK2* KO neurons, but not control neurons, 6 µM USP30i-37 (*Fig. 4B*) and 6 µM USP30i-3 (*Fig. 4C*) reduced TOM20 and HSP60 areas even in the absence of CCCP-induced mitochondrial damage, indicating that accumulated damaged mitochondria in the *PARK2* KO cells can be degraded in response to USP30 inhibition.

TEM analysis verified the TOM20 and HSP60 immunofluorescence data. In control neurons, but not *PARK2* KO neurons, mitochondrial abundance was reduced after CCCP treatment (*Figs. 5A-C*). USP30i-37 and USP30i-3 (3 µM) enhanced mitophagy in control and *PARK2* KO neurons subjected to mitochondrial damage by CCCP (*Figs. 5A-C*). Data also confirmed that USP30i-37 decreased the number of mitochondria in *PARK2* KO neurons but not in control neurons, even in the absence of CCCP-induced mitochondrial damage (*Figs. 5A-C*).

**Figure 5:**
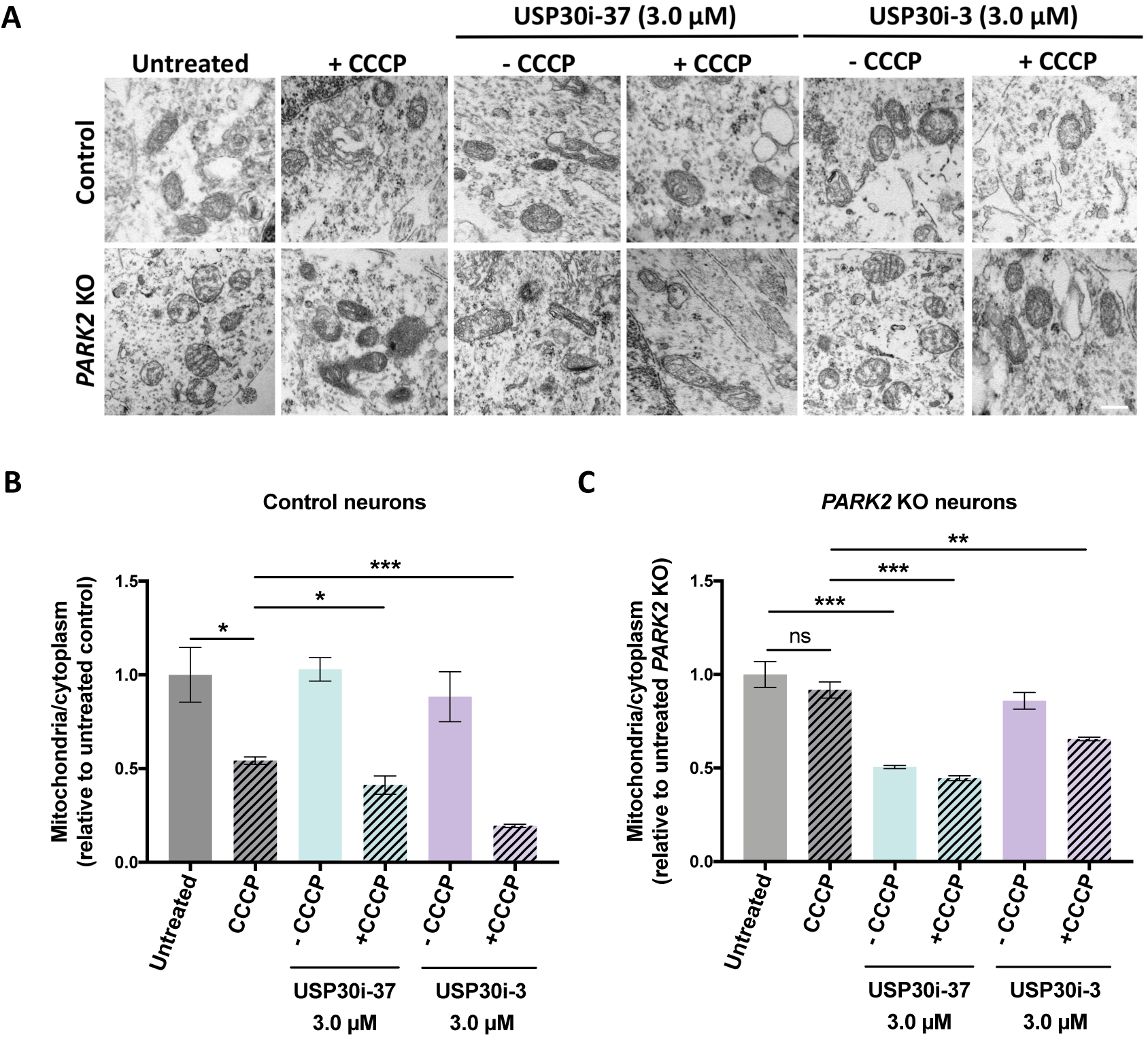
Transmission electron microscopy (TEM) confirms that USP30 inhibition induced mitophagy. **A)** TEM pictures showing mitochondria in control and *PARK2* KO neurons after treatment with 3 µM of USP30i-37 and USP30i-3 (+/- 10 µM of CCCP). Scale bar: 500 nm. Quantification of mitochondria/cytoplasm ratios showing the relative abundance of total and abnormal mitochondria in **B)** control and **C)** *PARK2* KO neurons. Data presented as mean±SEM, n=6 technical replicates, data from two independent differentiations. Significant differences are indicated by *p < 0.05, ***p < 0.001, ns: not significant, one-way ANOVA followed by Dunnett’s post hoc test for multiple comparisons.

Overall, USP30 inhibition exhibits the potential to boost mitophagy in CCCP-damaged control and *PARK2* KO neurons and to induce mitophagy at basal conditions in *PARK2* KO neurons.

### USP30 inhibition reduces excessive ROS production

ROS generation can cause oxidative damage to mitochondria, and the damaged mitochondria can promote further ROS generation, creating a vicious cycle that can cause cellular injury [32, 33]. Having shown that USP30 inhibition induces mitophagy, we investigated whether USP30 inhibition could reduce ROS levels. Differentiated neurons were treated with 0-6 µM USP30i-37 or USP30i-3 with or without CCCP, and hydrogen peroxide was measured at the cell population level as an indicator of oxidative stress. In control and *PARK2* KO neurons, ROS level significantly increased after CCCP exposure, indicating that CCCP treatment induces mitochondria-mediated oxidative stress (*Figs. 6A-D*). USP30i-37 (1.5, 3, and 6 µM) and USP30i-3 (3 and 6 µM in control; 1.5, 3, and 6 µM in *PARK2* KO) reduced CCCP-induced ROS levels in both cell lines (*Figs. 6A-D*). Interestingly, also basal ROS levels (no CCCP damage) in *PARK2* KO but not control neurons were reduced by 3 and 6 µM USP30i-37 and 6 µM USP30i-3 (*Figs. 6CD*), which may be linked to the higher basal ROS levels in the mutant line (*Fig. 2G*). This supports that mitophagy induction by USP30 inhibition can reduce ROS levels in compromised (either by CCCP or parkin KO) neurons.

**Figure 6:**
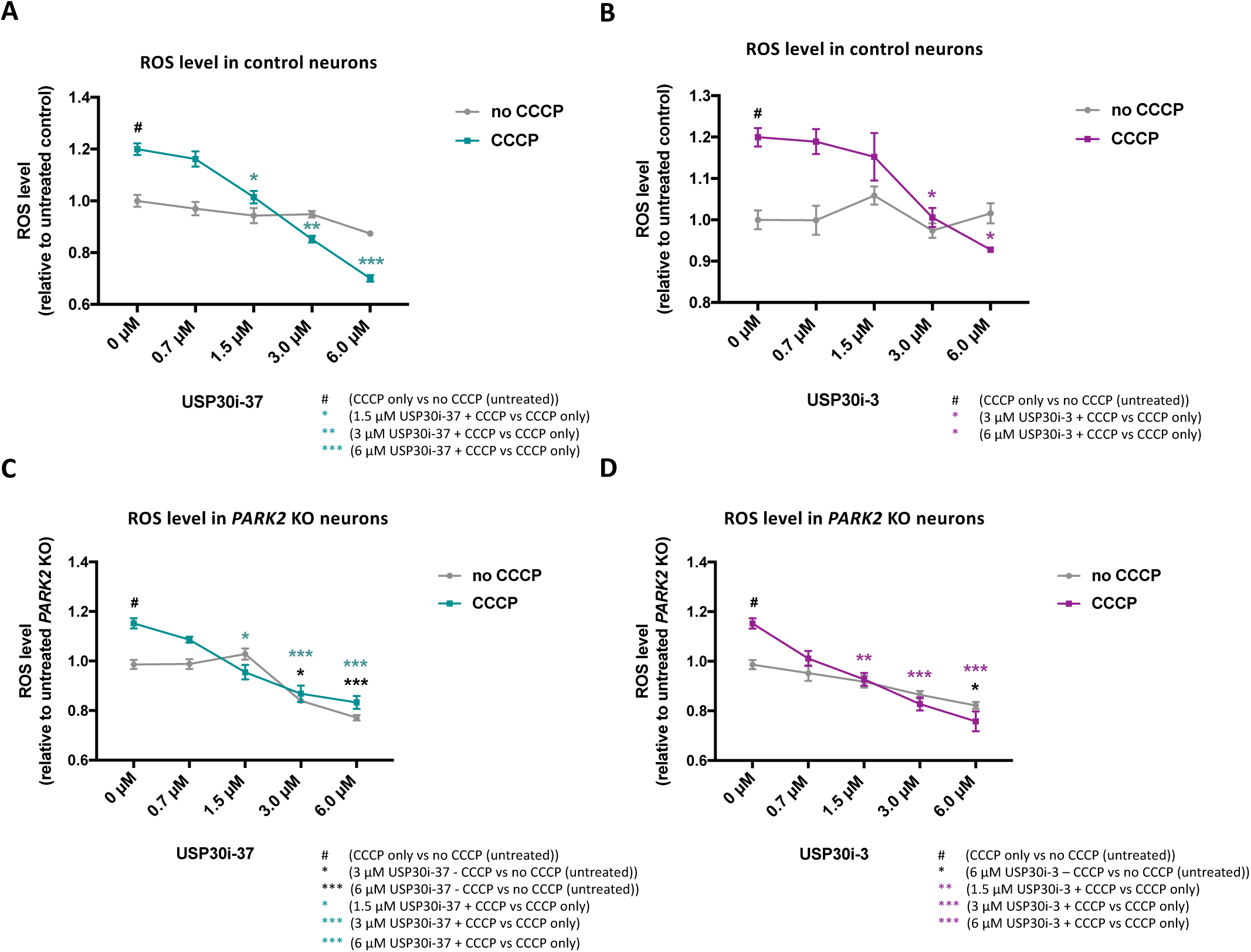
USP30 inhibition reduces excessive ROS production. Measurement of cellular H2O2 levels after USP30 inhibitors treatment (0.75 µM, 1.5 µM, 3 µM, 6 µM) added 4 h prior to CCCP (10 µM, 48 h) in **A)** control neurons treated with USP30i-37, **B)** control neurons treated with USP30i-3, **C)** *PARK2* KO neurons treated with USP30i-37, and **D)** *PARK2* KO neurons treated with USP30i-3. Data presented as mean±SEM, n=12 technical replicates, data from 3 independent differentiations. Significant differences are indicated by *, ^#^p < 0.05, **p < 0.01, ***p < 0.001, one-way ANOVA followed by Dunnett’s post hoc test for multiple comparisons.

### CCCP-induced mitophagy is impaired in both *PARK2* KO TH+ and TH-neurons, while USP30 inhibitor-induced mitophagy occurs mainly in TH-cells

Next, we investigated whether USP30 inhibitior-induced mitophagy was selective towards TH+ or TH-neuronal subtypes. *PARK2* KO neurons were exposed to 3 µM of USP30i-37 or USP30i-3, with or without CCCP (10 µM), and subsequently immunostained for HSP60 and TH. As demonstrated at the total population level (*Fig. 4*), CCCP did not reduce the HSP60 area in *PARK2* KO TH+ or TH-neurons (*Figs. 7A-C),* showing that mitophagy induction was impaired in both neuronal subtypes. The HSP60 area was reduced by 3 µM USP30i-3 but not USP30i-37 in both TH+ and TH-neurons after mitochondrial damage with CCCP (*Figs. 7A-C*). This is in line with USP30i-3 having a stronger effect than USP30i-37 at the total neuronal population level (*Fig. 4*). In TH-, but not TH+ neurons, 3 µM USP30i-37 and USP30i-3 reduced HSP60 areas in the absence of CCCP-induced mitochondrial damage, demonstrating a stronger effect of USP30 inhibition on basal mitophagy in the *PARK2* KO TH-neurons. It appears that TH-cells drive the USP30 inhibitior-mediated mitophagy in the total cell population, which is in accordance with the majority of neurons in the culture being TH-.

Effects of USP30 inhibition on oxidative stress were also investigated at the TH+ and TH-subpopulation level using the fluorogenic probe CELLROX, which exhibits bright green fluorescence upon oxidation by ROS and subsequent binding to DNA. The CCCP-induced ROS production was reduced by USP30i-37 and/or USP30i-3 in *PARK2* KO TH+ and TH-neurons (*Figs. 7D-F*). USP30 inhibition had no effect on basal ROS levels (no CCCP damage) in TH+ neurons (*Fig. 7E*) but reduced basal ROS levels in TH-neurons (*Fig. 7F*). This indicates that TH-neurons are the major drivers of the USP30 inhibitior-mediated reduction of basal ROS (no CCCP) in *PARK2* KO neurons (*Figs. 6CD*), supporting the observation that TH-neurons represent the treatment-sensitive population.

**Figure 7:**
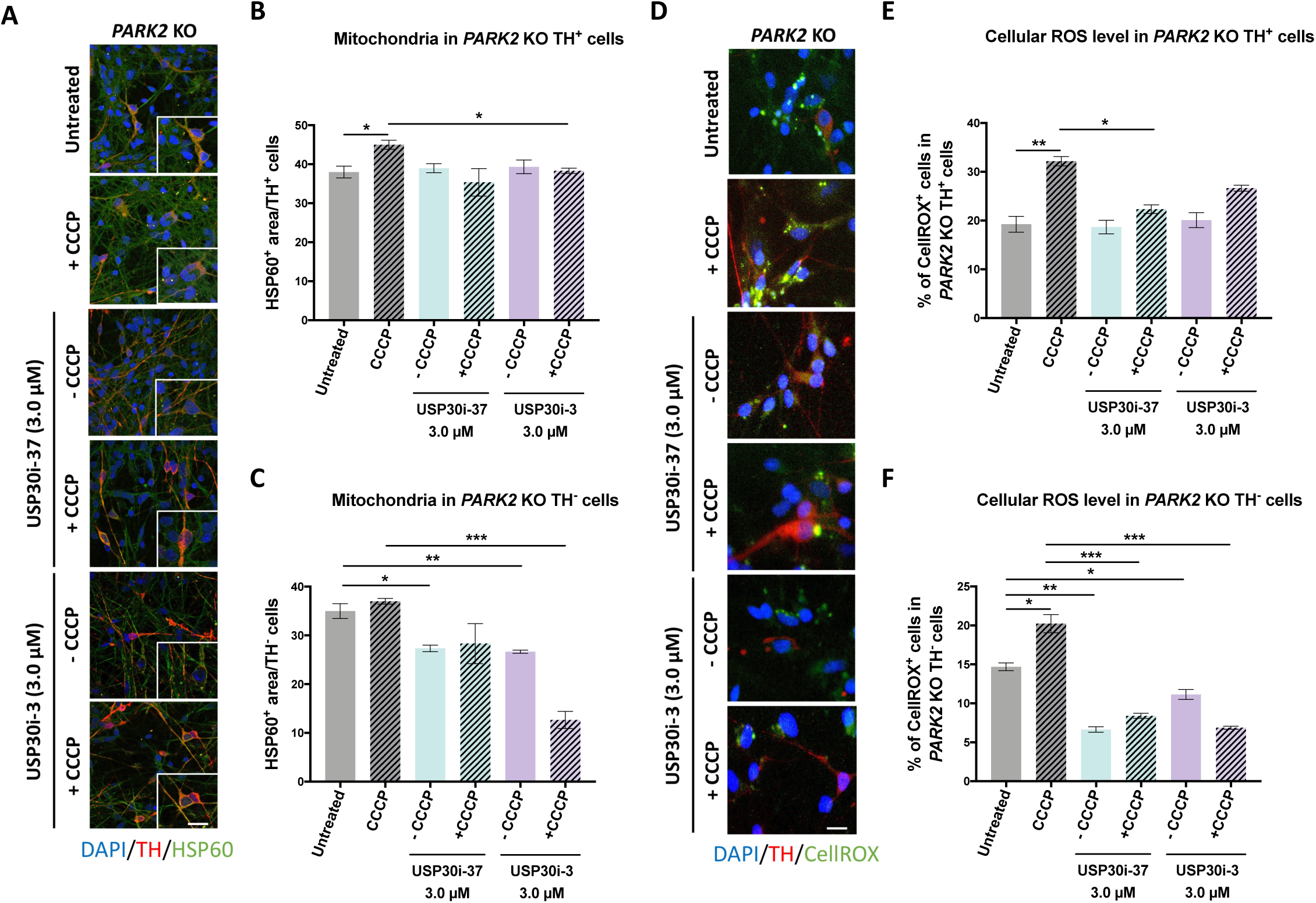
TH-negative neurons are the major driver of USP30 inhibitor-mediated mitophagy and ROS reduction in *PARK2* KO cells. **A)** Immunofluorescence staining for the mitochondrial marker HSP60 (green) and the catecholaminergic marker TH (red) after treatment with USP30i-37 and USP30i-3 (3 µM) added 4 h prior to CCCP (10 µM, 48 h). DAPI (blue) indicates nuclei. Scale bar: 50 µm. Quantification of HSP60 area upon USP30i-37 and USP30i-3 in **B)** *PARK2* KO TH+ neurons and **C)** *PARK2* KO TH-cells. **D)** Cellular ROS levels measured by CellROX Reagent (green) on catecholaminergic neurons (TH (red)) after treatment with USP30i-37 and USP30i-3 (3 µM) added 4 h prior to CCCP (10 µM, 48 h). DAPI (blue) indicates nuclei. Scale bar: 50 µm. Quantification of CellROX levels upon USP30i-37 and USP30i-3 treatment in **E)** *PARK2* KO TH+ neurons and **F)** *PARK2* KO TH-neurons. Data presented as mean±SEM, n=9 technical replicates from 3 independent differentiations. Significant differences are indicated by *p < 0.05, **p < 0.01, ***p < 0.001, one-way ANOVA followed by Dunnett’s post hoc test for multiple comparisons.

## Discussion

Accumulation of damaged mitochondria in parkin mutation carriers has been proposed to be the underlying pathogenesis of various diseases as E3 ligases add ubiquitin chains to the outer mitochondrial membrane proteins to trigger mitophagic removal of aged mitochondria [34]. We used our human iPSC-derived *PARK2* KO neuronal model to investigate the impact of parkin loss on mitochondrial turnover and function under basal conditions as well as under mitochondrial stress induced by treatment with the mitochondrial uncoupler CCCP. In line with our previous data [23–26], parkin deficiency resulted in increased mitochondrial area, accumulation of mitochondria with abnormal morphology, and increased ROS levels at basal conditions as compared to control neurons [25]. In the current study, we show that *PARK2* KO neurons are defective in clearing CCCP-impaired mitochondria. Several previous studies showed significantly impaired mitophagy in parkin-deficient cells [35–37], which correlates well with our observations of mitochondrial abnormalities and an impaired mitophagy response.

Both ubiquitin-dependent and -independent mitophagy pathways appear in the literature [18, 38]. It is unclear whether or not mitophagy in parkin mutation patients is driven by ubiquitination. Treatment with CCCP increased pS65-Ub levels in *PARK2* KO neurons, supporting the presence of ubiquitin labeling of mitochondria for degradation even in the absence of functional parkin, suggesting that another ubiquitin ligase must be in play. In fact, other E3 ubiquitin ligases such as HUWE1, MUL1, MARCH5 are reported to regulate mitophagy [39–41].

USP30 inhibition has been suggested as a therapeutic approach to facilitate ubiquitination-driven mitophagy in neurodegenerative diseases [13,42,43]. Our demonstration that mitochondria are labelled with ubiquitin following CCCP challenge in *PARK2* KO neurons is important as it highlights USP30 as a valid pharmacological target. As shown in this study, inhibition of USP30 appears to facilitate mitophagy. This is in line with the demonstration of pS65-Ub in *PARK2* KO mice and *PARK2* KO HEK cells [44, 45], but in contrast to the finding of negligible p-S65-Ub levels in human fibroblasts carrying heterozygote parkin mutations [18]. Differences may be due to the cell types, the sensitivity of the pS65-Ub assays, or the mitochondria decoupler treatment paradigms.

In our neuronal *in vitro* model, we found that USP30 inhibitors boosted CCCP-induced mitophagy in control neurons. This supports previous studies reporting enhanced CCCP-induced TOM20 and HSP60 reduction in USP30-depleted or USP30 KD cells [13,17,31]. Interestingly, our data indicate that USP30 inhibition facilitates mitophagy in parkin-deficient neurons at both basal growth conditions and upon CCCP-stress. In line with this, USP30 inhibition increases mitophagy in induced neurons derived from parkin loss-of-function carriers [46]. Moreover, defective mitophagy in SH-SY5Y cells transfected with PD-linked parkin mutants can be rescued with siRNA mediated knockdown of USP30 [13, 47]. In contrast, TOM20 reduction by USP30 inhibition requires parkin in HeLa cells [18].

We used our model system with *PARK2* KO or CCCP-induced mitochondrial impairment to investigate whether mitophagy induction by USP30 inhibition would lead to a healthier cellular environment. We found that parkin-deficient and/or CCCP-treated neurons generated more ROS than control cells and that USP30 inhibition reduced the level of oxidative stress. Overall, this indicates that USP30 inhibition can reduce ROS production, presumably by inducing mitophagy whereby damaged ROS-producing mitochondria are removed [48–50].

We show that USP30 inhibition induces mitophagy and reduces excessive ROS levels even in neurons that lack functional parkin, suggesting that USP30 also oppose parkin-independent mitophagy. CCCP-induced pS65-Ub levels were significantly higher in *PARK2* KO compared to control neurons, but *PARK2* KO neurons failed to induce mitophagy upon CCCP stress, pointing to the absence of mitophagy induction solely by the total ubiquitinate load as previously proposed [35]. One could speculate that the polyubiquitin chains linkage type, which could be different when other E3 ligases substitute for parkin, has a modulating effect on the potentiation of mitophagy [5].

Immunostaining for TH allowed us to investigate effects in TH+ and TH-neuronal subtypes in the dopaminergic differentiated cultures. While control neurons of both subtypes responded to CCCP damage with mitochondrial degradation, mitophagy was impaired in *PARK2* KO neurons of both subtypes. *PARK2* KO TH-neurons were more responsive to USP30 inhibition than TH+ neurons as they were the major driver of the USP30 inhibition-mediated mitophagy and ROS reduction. Although our data set does not indicate a strong beneficial effect of USP30 inhibition for TH+ neurons, it remains to be seen whether USP30 inhibition could improve dopaminergic cell survival in the intact brain, where cell-to-cell interaction is more diverse and complex than in the *in vitro* model.

In conclusion, our results support USP30 inhibition as a promising approach for promoting mitophagy of unhealthy mitochondria and reducing excessive cellular ROS production. Moreover, the approach seems valid in parkin-deficient cells. Although the *in vivo* effect on dopaminergic neurons in PD remains to be shown, the strategy could not only find applications in PD but also in other diseases with cellular dysfunction mediated by mitochondrial impairment.

## Acknowledgements

The authors would like to thank Dorte Lyholmer and Nadine Becker-von Buch for excellent technical assistance, Dr. Tore B. Stage for providing access to bioimaging equipment and Dr. Claire Gudex for editing the manuscript. The live imaging experiments reported in this paper were performed at DaMBIC, a bioimaging research core facility at the University of Southern Denmark. DaMBIC was established by an equipment grant from the Danish Agency for Science Technology and Innovation and by internal funding from the University of Southern Denmark. The research leading to these results was supported by H. Lundbeck A/S, the Innovation Fund Denmark (BrainStem & NeuroStem), the Danish Parkinson Foundation, the Jascha Foundation, the A.P. Møller Foundation for the Advancement of Medical Science (15-396, 14-427), and the Faculty of Health Sciences at the University of Southern Denmark.

## Author contributions

J.O., T.C.S., M.A., K.F., and M.M. designed research; J.O., J.B.A., T.C.S., and H.H. performed experiments; K.F and K.K.F. contributed new reagents or analytic tools; J.O., J.B.A., and H.H. analyzed data; J.O. and M.M. wrote the manuscript. T.C.S., M.A., K.K.F., and K.F. revised the manuscript for content. All authors read and approved the final manuscript.

## Material and methods

### Ethics approval

The Research Ethics Committee of the Region of Southern Denmark approved the study prior to initiation (S-20130101). All use of human stem cells was performed in accordance with the Danish national regulations and the ethical guidelines issued by the Network of European CNS Transplantation and Restoration (NECTAR) and the International Society for Stem Cell Research (ISSCR).

### *In vitro* propagation and differentiation of neural stem cells (NSCs)

Isogenic PARK2 KO and healthy control iPSC and NSC lines were provided by XCell Science Inc. (Novato, CA, USA). NSCs were propagated according to well-established, standard protocols using Geltrex-(Thermo Fisher)-coated plates in Neurobasal Medium (Thermo Fisher) supplemented with NEAA, GlutaMax-I, B27, supplement (Thermo Fisher), penicillin-streptomycin, and bFGF. Cells were enzymatically passaged with Accutase (Thermo Fisher) when 80-90% confluent. NSCs were differentiated according to a commercially available dopaminergic differentiation kit (XCell Science Inc.) for at least 25 days. Differentiation was divided into two parts: an induction phase, where NSCs were differentiated into dopaminergic precursors, and a maturation phase, where the dopaminergic precursor cells were differentiated into mature dopaminergic neurons. The differentiations were carried out at 37°C in a low, physiological O2 environment (5% CO2, 92% N2, and 3% O2). The cells were seeded onto wells coated with poly-L-ornithine (Sigma) and laminin (Thermo Fisher) at a density of 50,000 cells/cm^2^. Complete DOPA Induction Medium (XCell Science Inc.) supplemented with 200 ng/ml human recombinant Sonic Hedgehog (Peprotech) was changed every second day for the first nine days of differentiation. The cells were passaged at day 5 and day10 and seeded at a desired cell density. The medium was switched to Complete DOPA Maturation Medium A and B (XCell Science Inc.) at day 10 and 16, respectively. The generated neuronal populations were used for subsequent analyses.

### Treatments used

20 mM stock solution of *cyanide m-chlorophenylhydrazone* (CCCP, Sigma-Aldrich) was prepared by dissolving in dimethyl sulfoxide (DMSO) (Sigma-Aldrich) and further dilution to a final concentration of 10 µM. Cells were treated with CCCP in a time-dependent manner (24, 48, 72, and 96 h) to determine the optimal time; 48 h of exposure for further experiments was chosen.

USP30 inhibitors were derived from patent WO2016/156816 and synthesized in-house. USP30i-3 is the R-enantiomers of Example 3 and USP30i-37 is Example 37 in WO2016/156816. 100 mM stock solutions were prepared by dissolving in DMSO and further dilution in media to achieve final concentrations. USP30 inhibitor (0.75 µM, 1.5 µM, 3 µM, 6 µM) was added 4 h before addition of CCCP (10 µM).

All solutions were prepared on the day of the experiment. DMSO was used as a vehicle in all experiments and was included as untreated control (0 µM) (at 0.1% final concentration (v/v)).

### USP30 inhibitors concentration-response

To determine the highest non-toxic dose of the tested USP30 inhibitors, cells were differentiated as previously described for 36 days. At day 33 the cells were treated with 0 µM, 1.5 µM, 3 µM, 6 µM of USP30i-37 and USP30i-3 dissolved in DMSO. Cells treated with DMSO alone served as control. At day 36, cells were analyzed using lactate dehydrogenase (LDH) assay and stained for nuclei (DAPI) and neuronal markers (MAP2 and beta-TubulinIII).

### Assessment of cytotoxicity (LDH assay)

To assess cytotoxicity of the USP30 inhibitors in the tested doses, we used LDH assay to evaluate the content of necrotic cells. LDH assay was performed using CytoTox 96^®^Non-Radioactive Cytotoxicity Assay Kit (Promega) according to the manufacturer’s protocol. Briefly, cells were plated in 96-well plates (40,000 cells/well), and the absorbance was recorded at 490 nm using a microplate reader (VMAX kinetic ELISA).

### Western blotting

Western blotting was performed using standard techniques. In brief, the cells were lysed on ice in phosphate buffered saline (PBS) containing phosphatase (PhosphoSTOP Tablet, Roche) and protease (Complete Mini Tablets, Roche) inhibitors supplemented with 1% Triton X-100 (Sigma). Samples were then sonicated for 3x10 sec at amplitude 2 microns on ice, and the supernatants were collected. Protein content was measured with bicinchoninic acid assay (BCA, Pierce). Samples were equalized according to the protein concentration, denatured for 10 min in NuPAGE loading buffer, separated by SDS-PAGE on NuPage 4–12% Bis-Tris gel (NuPAGE) at 200V for 50 min with MOPS running buffer (NuPAGE). The proteins separated through iBlot transfer system (Invitrogen) were transferred onto polyvinylidene difluoride (PVDF) membranes (Invitrogen) followed by blocking the membranes for 60 min at 4°C with 5% non-fat dry milk in a mixture of 0.05 M Tris-Buffered Saline (TBS) and 0.05% Tween-20. The membranes were incubated overnight at 4°C in a solution containing the relevant primary antibodies. Subsequently, blots were incubated with an appropriate horseradish peroxidase (HRP)-conjugated secondary antibody for 1 h at RT. Protein expression was detected with ECL reagents (Thermo Fisher) using the ChemiDoc MP imaging system (BioRad) and quantified by densitometry using Image Lab software (BioRad).

*Primary antibodies:* mouse anti-Parkin (Cell Signalling #4211) 1:1000, mouse anti-USP30 (Abcam #AB3600) 1:500, rabbit anti-TOM20 (Santa Cruz #SC-11415) 1:1000, goat anti-HSP60 (Santa Cruz Biotechnology #SC-1052) 1:200, mouse anti-α-actin (Chemicon/Millipore #MAB1501) 1:6000.

*Secondary antibodies:* HRP-conjugated anti-mouse (DAKO #P0260) 1:2000, HRP-conjugated anti-rabbit (DAKO #P0790) 1:2000.

### Assessment of mitophagy

Cells were treated with 10 µM CCCP for 48 h to trigger mitophagy, and the process was evaluated by immunofluorescence staining for TOM20 (outer mitochondrial membrane marker) and HSP60 (mitochondrial matrix marker).

### Immunofluorescence staining

Cells cultured in 24- or 48-well plates (Corning) containing coverslips were fixed for 20 min at room temperature (RT) in 4% (w/v) paraformaldehyde (PFA, Sigma) in 0.15 M phosphate buffer (potassium dihydrogen phosphate (Merck) and disodium phosphate (Merck)), pH 7.4, and rinsed with 0.05 M TBS, pH 7.4, with 0.1% Triton-X-100 (Sigma). Cells were permeabilized, and unspecific binding was blocked with TBS containing 10% goat serum (Millipore) or donkey serum (Millipore) according to the host of the secondary antibodies. Cells were incubated overnight at 4°C with primary antibodies diluted in TBS/10% goat or donkey serum. Cultures were rinsed in TBS/0.1% Triton-X-100 and incubated with an appropriate secondary antibody diluted 1:500 in TBS/10% goat serum for 2 h at RT. Cell nuclei were counterstained with 10 μM 4",6-diamidino-2-phenylindole dihydrochloride (DAPI, Sigma) in TBS. Cultures were mounted onto glass slides with ProLong Diamond mounting medium (Molecular Probes).

*Primary antibodies:* mouse anti-β-III-tubulin (Sigma #T8660; 1:2000), mouse anti-microtubule-associated protein 2a+b (MAP2, Sigma #M1406; 1:2000), rabbit anti-tyrosine hydroxylase (TH, Millipore #AB152; 1:600), goat anti-forkhead box A2 (FOXA2 (Sigma #AF2400; 1:250), rabbit anti-GABA (Sigma #A2052; 1:2000), rabbit anti-GFAP (DAKO #Z0334; 1:4000), rabbit anti-TOM20 (Santa Cruz #SC-11415, 1:1000), goat anti-HSP60 (Santa Cruz Biotechnology #SC-1052, 1:200).

*Secondary antibodies:* Alexa Fluor 555 goat anti-mouse IgG (Molecular Probes #A21422, 1:500), Alexa Fluor 594 donkey anti-goat (ThermoFisher #A11058, 1:500), Alexa Fluor 488 goat anti-rabbit IgG (ThermoFisher #A11008, 1:500), Alexa Fluor 488 donkey anti-goat IgG (Invitrogen #A11055, 1:500).

### Image analysis

Fluorescence pictures were taken using a FluoView FV1000MPE – Multiphoton Laser Confocal Microscope (Olympus) 60X magnification, in a blinded manner on 5 randomly chosen confocal fields per coverslip from independent experiments. Mitophagy was analyzed by quantifying the total area of TOM20+ and HSP60+ stainings. The analysis was performed automatically in ImageJ Software by converting images to binary format and analyzing particles. The total area of TOM20+ and HSP60+ stainings was normalized to the total cell numbers as quantified automatically in ImageJ for Hoechst+/DAPI+ nuclei.

### Transmission electron microscopy (TEM)

Cells were seeded on 13 mm Thermanox plastic coverslips (Nunc) coated with poly-L-ornithin/laminin, primarily fixed in 3% glutaraldehyde (Merck) in 0.1 M sodium phosphate buffer with pH 7.2 at 4°C for 1 h, and stored in a 0.1 M Na-phosphate buffer at 4°C until further analysis. When ready, the cells were embedded in 4% agar at 45°C (Sigma) under the stereomicroscope and cut into 1-2 mm^3^ blocks, which were then washed with 0.1 M Na-phosphate buffer followed by post-fixation in 1% osmium tetroxide in 0.1 M Na-phosphate buffer (pH 7.2) for 1 h at RT. Cells were washed in MilliQ water, followed by stepwise dehydration in a series of ascending ethanol concentrations ranging from 50-99% EtOH. Propylene oxide (Merck) was then used as an intermediate to allow infiltration with Epon (812 Resin, TAAB). The following day, the agar blocks were placed in flat molds in pure Epon, cured at 60°C for 24 h. Approximately eight semi-thin sections (2 μm) from one block were cut on an ultramicrotome with a glass knife (Leica, Reichert Ultracut UTC). These were stained with 1% toluidine blue in 1% Borax and evaluated by light microscopy to locate areas with adequate number of cells for further processing. Ultra-thin sections (70 nm) were cut on the ultramicrotome with a diamond knife (Jumdi, 2 mm) and then collected onto TEM copper grids (Gilder) and stained with 2% uranyl acetate (Polyscience) and 1% lead citrate (Reynolds 1963). The samples were evaluated, and the images were collected using a Philips CM100 transmission electron microscope equipped with a Morada digital camera equipment and iTEM software system.

### Morphometric analysis of TEM images

Six TEM grids from each cell line were used for analysis, and ten spots were randomly chosen at low magnification. Images of each spot were captured using high magnification (19,000X). To estimate mitochondrial abundance, the organelles were counted manually on each micrograph and normalized to cytoplasm area. Thirty images were used for each cell line.

### Phospho-Ubiqutin (Ser65) assay

Phosporylated Ubiqutin at Ser65 levels were measured using time-resolved fluorescence resonance energy transfer (FRET) Phospo-Ubiqutin (Ser65) Cellular Kit (Cisbio 64UBIS65PEG) according to the manufacturer’s instructions. Briefly, cells were plated in a 96-well plate (60,000 cells/well) and lysed using lysis buffer for the kit. Cell lysate samples were transferred into a new plate, and Activation Buffer was added together with Mix Detection Antibodies. The plate was incubated for 1 h at RT and read on an HTRF plate reader (Molecular Devices).

### Measurement of oxidative stress levels

#### ROS assay

ROS level was measured using ROS-Glo™ H2O2 Assay Kit (Promega) according to the manufacturer’s instructions. Briefly, cells were plated in a 96-well plate (5,000 cells/well). Media were transferred to a new 96-well plate, and the H2O2 substrate was added to each media sample and incubated for 1 h at 37°C. The ROS-Glo™ Detection Solution was then added, the plate was incubated for additional 20 min at RT, and the luminescence signal was recorded using an Orion L Microplate Luminometer (Titertek Berthold).

#### CellROX assay

Cellular ROS levels were measured using CellROX™ Green Reagent (ThermoFisher, Invitrogen) according to the manufacturer’s instructions. Briefly, cells were plated in a 96-well plate (40,000 cells/well). The CellROX Reagent was added to the cells at a final concentration of 5 µM, and the plate was incubated for 30 min at 37°C. Medium was removed, and the cells were washed 3 times with PBS. Cells were fixed with 4% PFS for 10 min and stained for DAPI and TH. Images were generated within 24 hours using ImageIxpress Pico Automated Cell Imaging System (Molecular Devices). Image analysis was performed by quantifying the staining area and normalizing to DAPI+nuceli using ImageJ program.

### Statistical analysis

Statistical analysis was performed using GraphPad Prism 7.0 software. We applied two-tailed unpaired Student’s t-test and one- or two-way ANOVA with multiple comparison test. Data were considered statistically significant at p < 0.05 (*, ^#^), p < 0.01 (**, ^##^), and p < 0.001 (***, ^###^). Data are presented as mean ± standard error of the mean (SEM).

## Figure titles and legends

**Suppl. Figure S1:**
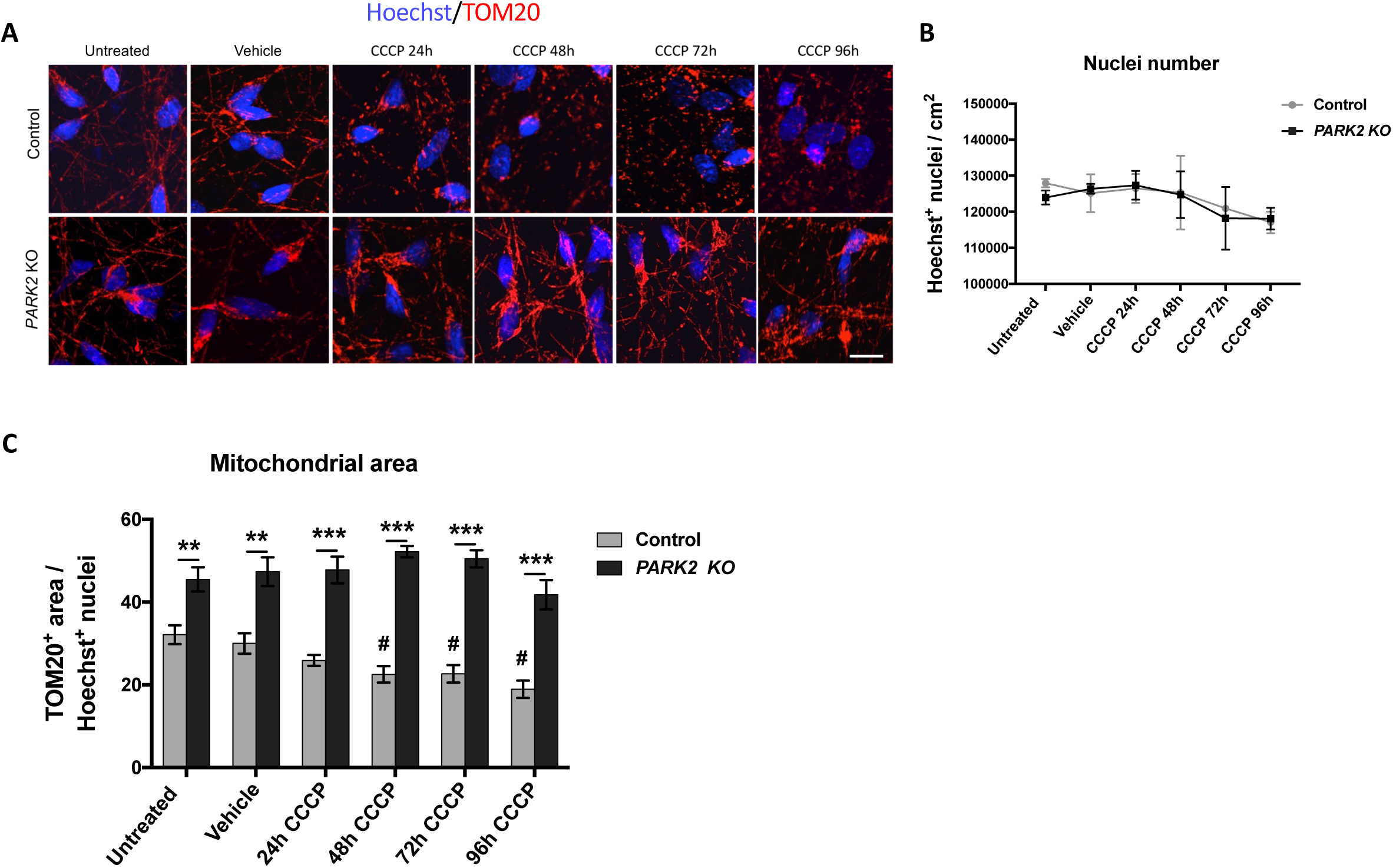
Impaired CCCP-induced TOM20 degradation in *PARK2* KO neurons. **A)** *PARK2* KO and control neurons were untreated or treated with vehicle (DMSO) or 10 μM CCCP for 24, 48, 72, and 96 hrs and immunofluorescence stained for TOM20 (red) and Hoechst (blue) to visualize mitochondria and nuclei as indicated. Scale bar: 10 μm. **B)** No significant differences in total cell numbers were observed upon CCCP or vehicle (0.1% DMSO) exposure although there was a clear trend towards fewer cells after 72 and 96 hrs. **C)** Quantification of TOM20+ mitochondrial area normalized to number of Hoechst+ nuclei showed that the mitochondrial area in control neurons was significantly reduced after 48, 72, and 96 hrs of CCCP treatment. In contrast, mitochondria were retained in the *PARK2* KO neurons. Vehicle (0.1% DMSO) did not affect area of TOM20 immunoreactivity. Data presented as mean±SEM, n=9 technical replicates, data from 3 independent differentiations, Significant differences are indicated by **p < 0.01, ***p < 0.001 (control vs. *PARK2* KO), ^#^p < 0.05 (untreated control vs. CCCP-treated control), one-way ANOVA followed by Dunnett’s post hoc test for multiple comparisons

**Suppl. Figure S2:**
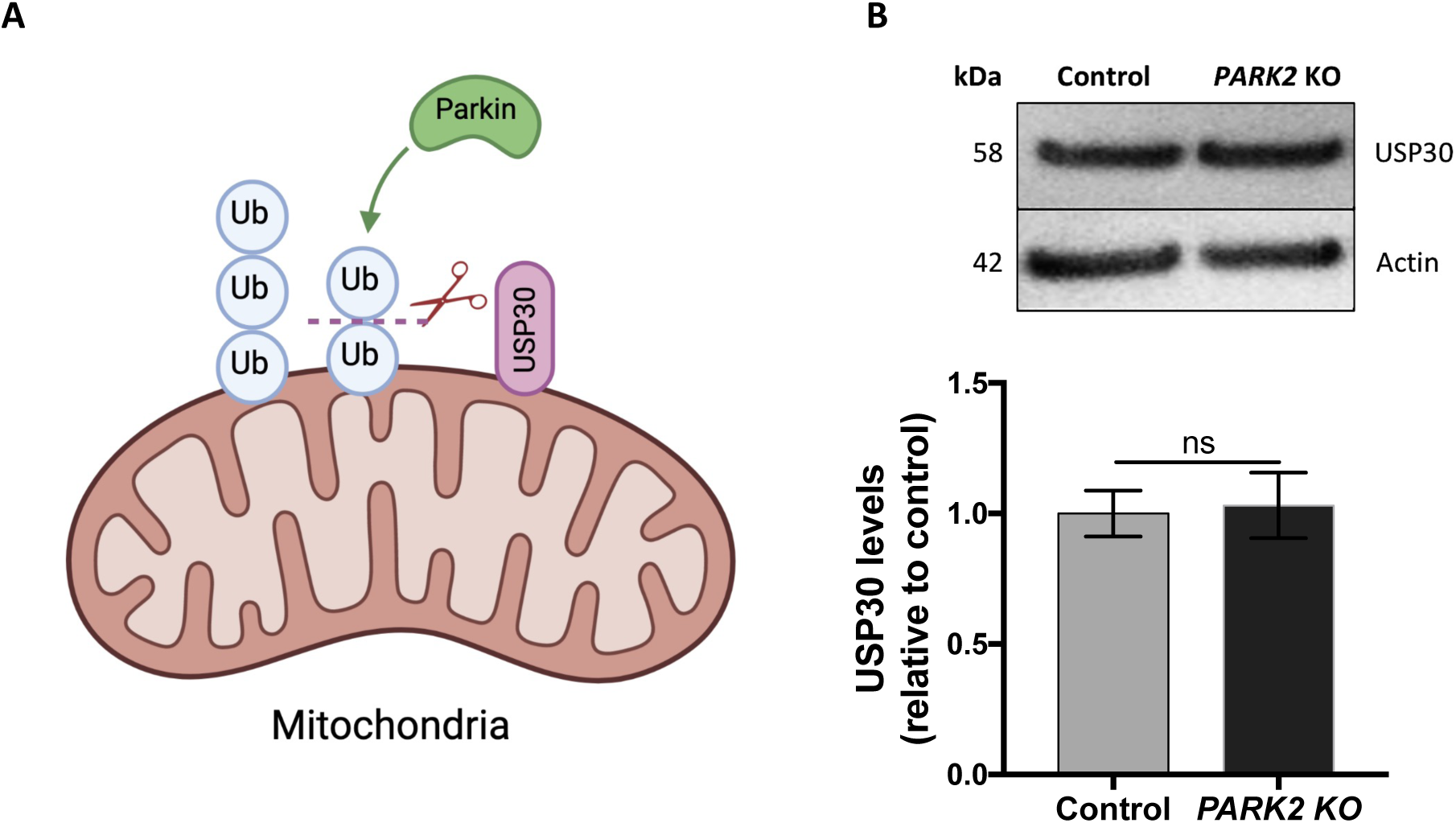
USP30 inhibition as a strategy to enhance mitophagy. **A)** USP30 opposes parkin-mediated ubiquitination. **B)** Western blotting and densitometric analysis demonstrating comparable levels of USP30 protein expression in *PARK2* KO and control neurons. Representative blots of two independent experiments.

**Suppl. Figure S3:**
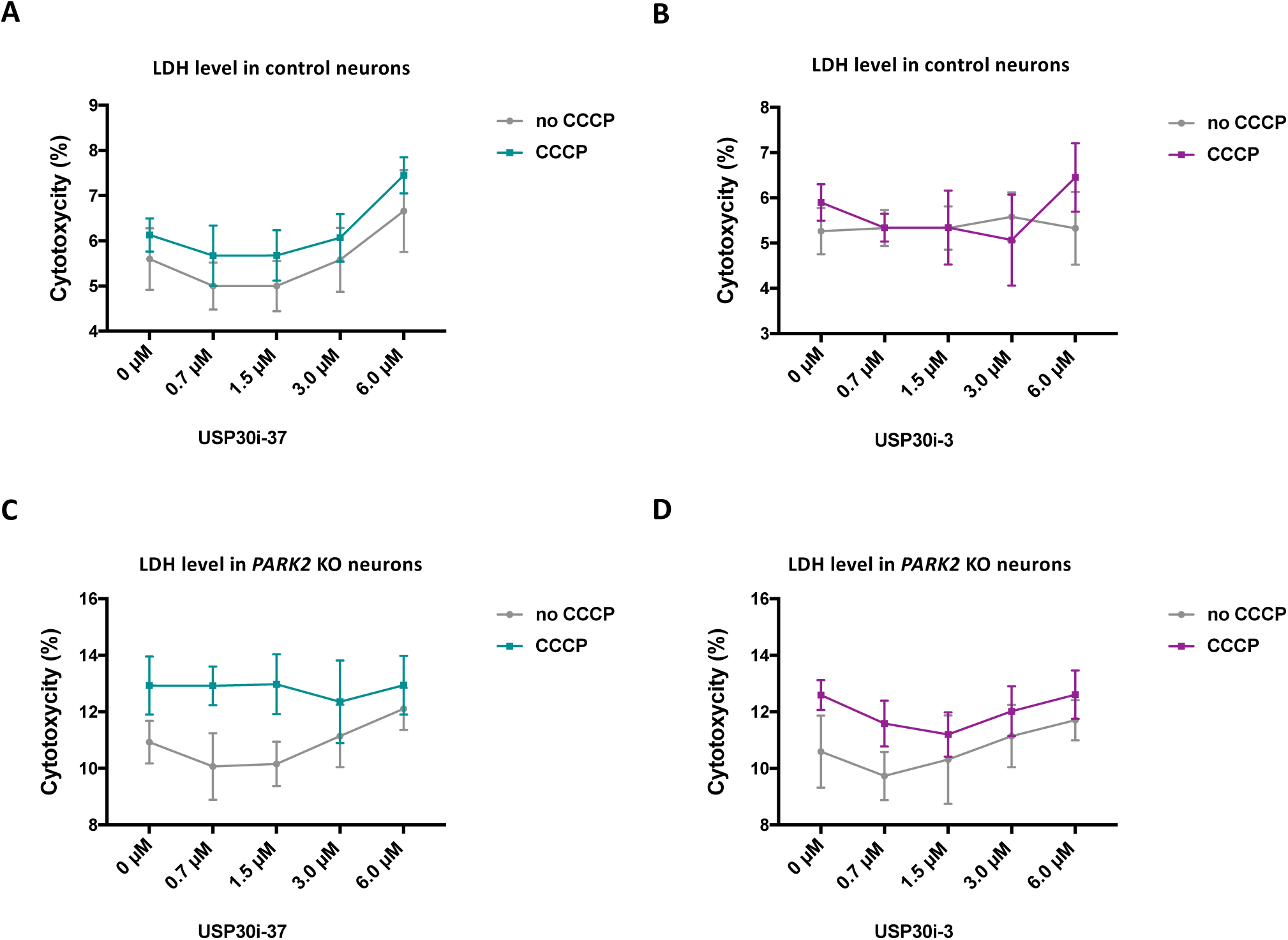
Lactate dehydrogenase (LDH) release did not reveal cytotoxic effects of USP30 inhibitors (+/-CCCP) Differentiated neurons were treated with with 0 µM (untreated), 0.75 µM, 1.5 µM, 3 µM, 6 µM of USP30i-37 and USP30i-3 inhibitors added 4 h prior to CCCP (10 µM, 48 h). Necrotic cell death was assessed by LDH analysis of conditioned media from **A)** control neurons treated with USP30i-37, **B)** control neurons treated with USP30i-3, **C)** *PARK2* KO neurons treated with USP30i-37, and **D)** *PARK2* KO neurons treated with USP30i-3. No significant cytotoxicity was observed after treatment in either cell line, but there was a tendency to increase with increasing dose. Data presented as mean ± SEM, n=9-15 technical replicates, data from 3 independent differentiations.

**Suppl. Figure S4:**
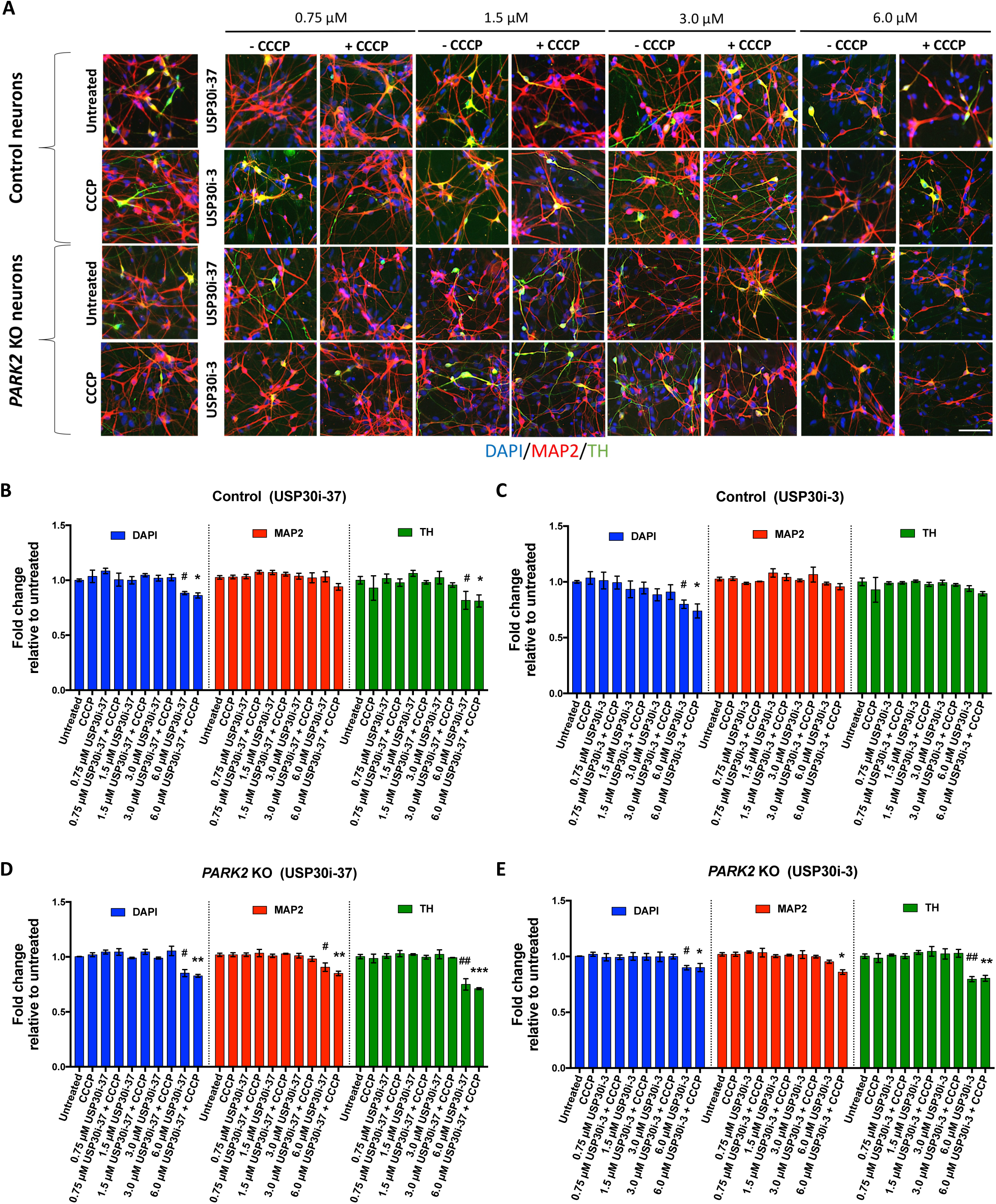
Immunofluorescence staining showed that 3 µM USP30 inhibitors did not affect the numbers of nuclei (DAPI), mature neurons (MAP2), or dopaminergic neurons (TH) (+/-CCCP) Healthy control and *PARK2* KO neurons were treated with 0 µM (untreated), 0.75 µM, 1.5 µM, 3 µM, 6 µM of USP30i-37 and USP30i-3. Cells were fixed and stained for DAPI (blue), MAP2 (red), and TH (green). **A)** Representative immunoflorescence pictures of DAPI+, MAP2+, and TH+ cells in healthy control and *PARK2* KO iPSC-derived neurons. Scale bars: 200 µm. **B-E)** Quantification of total cell count in **B, C)** healthy control and **D, E)** *PARK2* KO neurons. The toxicity of USP30 compounds (decreased cell number and compromised cell morphology) increased in both cell lines at the highest concentrations. Data presented as mean±SEM, n=9 technical replicates, data from 3 independent differentiations. Significant differences are indicated by *, ^#^p < 0.05, **, ^##^p < 0.01, ***p < 0.001, one-way ANOVA followed by Dunnett’s post hoc test for multiple comparisons.

